# Characterization of a set of abdominal neuroendocrine cells that regulate stress physiology using colocalized diuretic peptides in *Drosophila*

**DOI:** 10.1101/164178

**Authors:** Meet Zandawala, Richard Marley, Shireen A. Davies, Dick R. Nässel

## Abstract

Multiple neuropeptides are known to regulate water and ion balance in *Drosophila melanogaster*. Several of these peptides also have other functions in physiology and behavior. Examples are corticotropin-releasing factor-like diuretic hormone (diuretic hormone 44; DH44) and leucokinin (LK), both of which induce fluid secretion by Malpighian tubules (MTs), but also regulate stress responses, feeding, circadian activity and other behaviors. Here, we investigated the functional relations between the LK and DH44 signaling systems. DH44 and LK peptides are only colocalized in a set of abdominal neurosecretory cells (ABLKs). Targeted knockdown of each of these peptides in ABLKs leads to increased resistance to desiccation, starvation and ionic stress. Food ingestion is diminished by knockdown of DH44, but not LK, and water retention is increased by LK knockdown only. Thus, the two colocalized peptides display similar systemic actions, but differ with respect to regulation of feeding and body water retention. We also demonstrated that DH44 and LK have additive effects on fluid secretion by MTs. It is likely that the colocalized peptides are coreleased from ABLKs into the circulation and act on the tubules where they target different cell types and signaling systems to regulate diuresis and stress tolerance. Additional targets seem to be specific for each of the two peptides and subserve regulation of feeding and water retention. Our data suggest that the ABLKs and hormonal actions are sufficient for many of the known DH44 and LK functions, and that the remaining neurons in the CNS play other functional roles.

## Introduction

Orchestration of physiological and behavioral processes is commonly dependent on neuropeptide and peptide hormone signaling [see 1,2–4]. For example, feeding and postprandial effects on the organism, including satiety, nutrient and energy reallocation, diuresis, and activity/sleep are regulated by multiple peptides [see 5,6,1,7–9,3]. Thus, in *Drosophila melanogaster*, several neuropeptides such as allatostatin A, neuropeptide F, short neuropeptide F (sNPF), sulfakinin, and hugin-pyrokinin are known to regulate feeding, and five peptides, diuretic hormones 31 and 44, leucokinin (LK) as well as CAPA-1 and 2, derived from the gene *capability*, regulate ion and water homeostasis [see 10,11,12,1,7,3,13]. After a meal other hormones, like insulin-like peptides (ILPs) and adipokinetic hormone (AKH) ensure energy mobilization and storage, or signal satiety/hunger and affect organismal activity, vigor and stress tolerance [14–16,7].

The neuroendocrine cells producing the peptides mentioned above display varying degrees of diversity, from a single small set of identical cells producing AKH or ILPs to very large populations of diverse neurons expressing sNPF [17–19]. Thus, the question is whether peptidergic neurons of a large diverse population are functionally coupled and play a concerted physiological role, or if they are parts of distributed networks where the specific neuropeptide therefore serve diverse functions. To address this question we have selected a set of neuroendocrine cells producing the neuropeptide LK that consists of four morphological types of cells [20,21], and is proposed to serve multiple functions in flies [22–26,6]. This set of neurons was dissected by genetic tools to enable us to isolate the functional role of a major subset, which consists of prominent neurosecretory cells in the abdominal ganglia. We show that this subset of the LK neurons, designated ABLKs, additionally produce corticotrophin-releasing factor-like diuretic hormone, also known as diuretic hormone 44 (DH44). These ABLKs are especially intriguing since they seem to be under tight neuronal and hormonal control. Receptors for several neurotransmitters and peptides have been identified on these cells in larvae [27,28,24,29–32] and adults [24]. Of these receptors, only the action of the 5-HT1B receptor on ABLK function was probed in adults [24]. We ask what function these specific neuroendocrine cells and their colocalized peptide hormones have in physiology and behavior.

Both DH44 and LK are primarily known for their roles as diuretic hormones in various insects, including *Drosophila*, by regulating secretion by Malpighian (renal) tubules (MTs) [33,12,3,34]. However, several additional functions have been assigned to these peptides from genetic experiments. LK neurons regulate food intake, play a role in desiccation stress resistance, modulate chemosensory responses, decrease postprandial sleep, and are required for starvation-induced sleep suppression [22–26,6]. DH44, which is produced by a diverse set of neurons and neurosecretory cells [33,35], plays roles in osmotic and metabolic stresses. Knockdown of DH44 in the CNS or its receptor (DH44-R2) in the MTs results in a significant increase in desiccation tolerance [23]. Genetic ablation of DH44 neurons also results in increased starvation tolerance, however, knockdown of the DH44 receptor, DH44-R2, in the MTs impairs starvation tolerance [23]. Furthermore, DH44 producing median neurosecretory cells in the brain regulate rhythmic locomotor activity with influence from clock cells [36], sense and regulate intake of nutritive carbohydrates [37], and regulate sperm retention in the uterus of females [35].

The postulated functions of LK and DH44 are, with a few exceptions, not assigned to specific neurons. Using the GAL4-UAS system [38] we interfered with LK and DH44 expression in the ABLKs and analyzed *in vivo* effects on tolerance to various stressors, as well as feeding and water retention. We also employed an assay to monitor the combined activity of DH44 and LK on secretion in MTs. These peptides act on different cell types of the *Drosophila* MTs and activate different signaling pathways [33,34], yet we show that they display an additive stimulatory effect on secretion. Thus we can show that the ABLKs, and therefore, the hormonal actions of the two peptides, are sufficient for regulating water and ion homeostasis and associated stress functions, but can also affect food intake, perhaps by an indirect action caused by diuretic activity. This suggests that the LK and DH44 neurons in the brain are important for the additional functional roles listed above, and it remains to be determined whether these functions are in any way linked to those of the ABLKs.

## Experimental procedures

### Fly lines and husbandry

All fly strains used in this study (Table 1) were reared and maintained at 25°C on a standard yeast, corn meal, agar medium (see http://flystocks.bio.indiana.edu/Fly_Work/media-recipes/bloomfood.htm) supplemented with 1.5 g/L nipagin and 3 mL/L propionic acid. Experimental flies were reared under uncrowded conditions and normal photoperiod (12 hours light: 12 hours dark).

**Table 1:** Fly strains used in this study

| Fly strain | Inserted on chromosome | Source / reference | Stock number |
| --- | --- | --- | --- |
| *w<sup>1118</sup>* | - | BDSC | - |
| *w<sup>1118</sup>; Lk-GAL4* | II | [21], Pilar Herrera, Madrid, Spain | - |
| *w<sup>1118</sup>; Lk-GAL4* | II | [39] | BDSC 51993 |
| *w<sup>1118</sup>; Lk-GAL4* | III | Young Joon Kim, Gwangju, Korea | - |
| *w<sup>1118</sup>; DH44-GAL4* | III | [40], Fly Light | BDSC 39347 |
| *w<sup>1118</sup>; DH44-GAL4* | III | [39] | BDSC 51987 |
| *y<sup>1</sup>w<sup>\*</sup>; Actin5c-GAL4/CyO* | II | - | BDSC 4414 |
| *w<sup>1118</sup>; UAS-Lk RNAi* | III | [41] | VDRC<br>14091/GD |
| *yw; UAS-DH44 RNAi* | II | - | VDRC<br>108473/KK |
| *yw; Sco/CyO; UAS-mcd8-GFP* | III | BDSC | - |
| *JFRC29-10xUAS-IVS-myr::GFP-p10* | - | [42], Washington, USA | - |

### Immunohistochemistry and imaging

Immunohistochemistry for *Drosophila* larval and adult CNS was performed as described earlier [43]. Briefly, CNS from third instar larvae or adult male flies were dissected in phosphate buffered saline (PBS). Larval samples were fixed for 2 hours in 5% ice-cold paraformaldehyde and adult samples were fixed on ice for 3.5 – 4 hours. Samples were then washed in PBS and incubated for 48 hours at 4°C in primary antibodies diluted in PBS with 0.5% Triton X (PBST) (Table 2). Following this incubation, samples were washed with PBST and incubated for 48 hours at 4°C in secondary antibodies diluted in PBST (Table 2). Next, all samples were thoroughly washed with PBST, and following a final wash in PBS, samples were mounted in 80% glycerol. For anti-DH44 staining, tissues were blocked with 5% normal goat serum (NGS) in PBST post-fixation and 5% NGS was also included in the primary antibody solution.

**Table 2:** Antibodies used for immunohistochemistry

| Antibody | Immunogen | Source / reference | Dilution |
| --- | --- | --- | --- |
| **Primary antibody** |  |  |  |
| Rabbit anti-LK | *Leucophaea maderae* kinin I | [44] | 1:2000 |
| Rabbit anti-DH44 | *Drosophila melanogaster* DH44 | [33] Jan Veenstra, Bordeaux, France | 1:1000 |
| Mouse anti-GFP | Jelly fish GFP | Invitrogen | 1: 1000 |
| **Secondary antibody** |  |  |  |
| Goat anti-mouse Alexa 488 | - | Invitrogen | 1:1000 |
| Goat anti-rabbit Alexa 546 | - | Invitrogen | 1:1000 |

All samples were imaged with a Zeiss LSM 780 confocal microscope (Jena, Germany) using 10X, 20X or 40X oil immersion objectives. Confocal images were processed with Zeiss LSM software and Fiji [45] for projection of z-stacks, contrast and brightness, and calculation of immunofluorescence levels. Cell fluorescence was measured as described previously [43]. Briefly, the cells of interest were selected and their area, integrated density and mean gray values measured. Background values for these parameters were also recorded by selecting a region that has no fluorescence near the cells of interest. Corrected total cell fluorescence (CTCF) was then calculated using the equation: CTCF = Integrated Density – (Area of selected cell X Mean fluorescence of background readings).

### Stress resistance assays

We used 5-6 days old male flies to assay for survival under various stresses and recovery from chill coma (see [43] for details of stress assays). For each technical replicate, 15 flies were kept in a vial and their survival recorded every 3 hours (for desiccation) or 6 hours (for starvation and ionic stress) until all the flies were dead. For desiccation, flies were kept in empty vials. For starvation, flies were kept in vials containing 5 ml of 0.5% aqueous agarose (A2929, Sigma-Aldrich). For ionic stress, flies were kept in vials containing 5 ml enriched medium (100 g/L sucrose, 50 g/L yeast, 12 g/L agar, 3 ml/L propionic acid and 3 g/L nipagin) supplemented with 4% NaCl. All vials were kept at 25°C under normal photoperiod conditions for the entire duration of the experiment. For chill coma recovery experiments, flies were transferred to empty vials, which were then placed on ice to induce a chill coma. The vials were incubated on ice (0°C) for 4 hours and afterwards transferred to 25°C to induce recovery. The number of flies recovered was assessed every 2 min. At least 3 biological replicates and 3 technical replicates for each biological replicate were performed for each experiment.

### Capillary feeding assay

Capillary feeding (CAFE) assay to measure food intake for individual flies was performed according to the method described earlier [24]. Food consumption was measured daily and the cumulative food intake over 4 days was calculated. The experiment consisted of 3 biological replicates and 8-10 flies per replicate for each genotype.

### Water content measurement

For measurement of water content, 10-15 flies were frozen on dry ice and their weight (wet weight) recorded using a Mettler Toldeo MT5 microbalance (Columbus, USA). The flies were then dried for 1 day at 60°C and their weight (dry weight) recorded again. The water content of the flies was determined by subtracting dry weight from wet weight (see [46]).

### Malpighian tubule secretion assay

*Drosophila* MT fluid secretion assays were performed as described previously [47]. Briefly, MTs from six day old *Drosophila* were dissected in Schneider’s medium and transferred to 9 μl of 50% Schneider’s medium and 50% *Drosophila* saline [33]. Tubules were left to secret for 30 minutes and non-secreting tubules were replaced to form a data set of 10-15 secreting tubules. Basal secretion was measured for 30 min at 10 min intervals. Following this initial incubation, 1 μl of *Drosophila* DH44 (final concentration 10^−7^ M or 10^−9^ M), LK (final concentration 10^−9^ M or 10^−10^ M) or both (Genosphere Biotechnologies, Paris, France) were added to the incubation medium. Stimulated secretion was measured for 30 min at 10 min intervals. Data are presented as the secretion rate at every time point, the percentage change in secretion rate following peptide application, and total fluid secreted over 60 min.

### Statistical analyses

The experimental data are presented as means ± s.e.m. Unless stated otherwise, oneway analysis of variance (ANOVA) followed by Tukey’s multiple comparisons test was used for comparisons between three genotypes and an unpaired t-test was used for comparisons between two genotypes. For fluid secretion assays, a Mann-Whitney U test was used as some data were non-normally distributed. Stress curves were compared using Mantel-Cox log-rank test. All statistical analyses were performed using GraphPad Prism with a 95% confidence limit (*p* < 0.05).

## Results

### LK expression in Drosophila CNS

Several studies have previously examined the distribution of LK in *Drosophila* CNS and peripheral tissues [20,21,24,48,23]. Here, we verified the expression of *Lk-GAL4* driven GFP in both larval (Figure S1A-E) and adult *Drosophila* (Figure 1), using a GAL4 line from <u>De Haro et al.</u> [21]. In the larval CNS, *Lk-GAL4* drives expression in five pairs of neurons in the brain (Figure S1A, E), three pairs of neurons in the SOG (Figure S1C, E) and seven pairs of neurons in the VNC (Figure S1D, E). However, four out of the five pairs in the brain do no display any LK-immunoreactivity, but in fact react with an antiserum to ion transport peptide (ITP) (Figure S1B). These lateral neurosecretory cells are found both in larval and adult brains and are known as ipc-1 neurons [49]. Similar to the larval CNS, *Lk-GAL4* drives GFP expression in four distinct neuronal populations in the adult CNS (Figure 1). Hence, GFP expression was detected in one pair of neurons in the lateral horn and one pair in the SOG (Figure 1A, D), another four pairs in the brain, which display ITP-immunoreactivity (Figure 1B, D and S2) and eleven pairs in the VNC (Figure 1C, D). In adult flies, *Lk-GAL4* driven GFP expression in the ITP-producing cells is weak and variable (Figure S2). Interestingly, seven pairs of neurons in the VNC start expressing LK in the embryonic stage, whereas the additional four pairs (this number varies between individuals) begin to express LK during pupal development [48].

Moreover, *Lk-GAL4* also drives ectopic expression in salivary glands (Figure 1E). Although we tested two additional *Lk-GAL4* driver lines (Table 1), we utilized this *Lk-GAL4* for subsequent knockdown experiments because its expression is stronger than, or more specific than the other *Lk-GAL4* lines (Table 1; data not shown).

**Figure 1:**
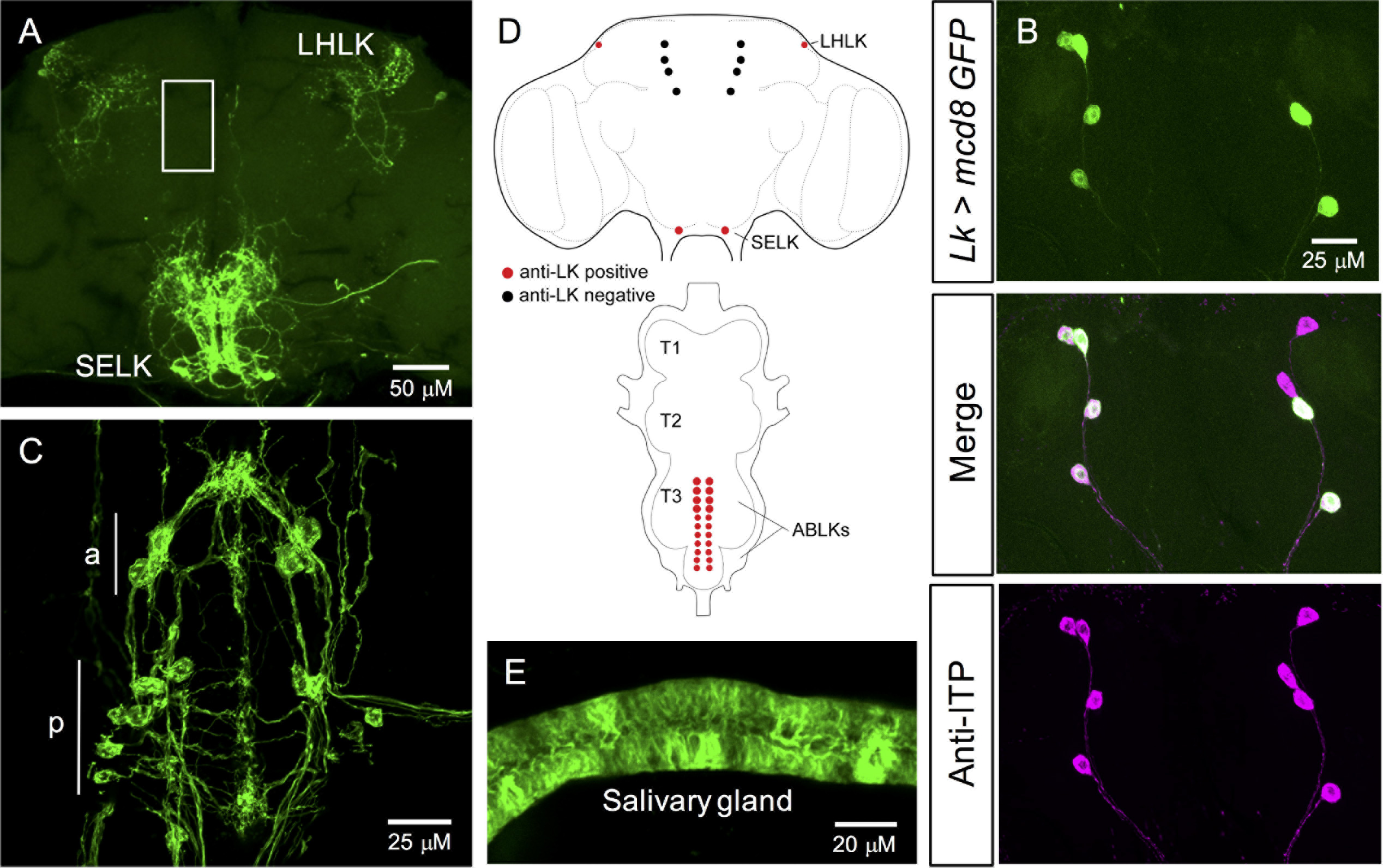
The *Lk-GAL4* drives GFP expression in four distinct neuronal populations in the adult *Drosophila* CNS. **(A)** One pair of neurons in the lateral horn (lateral horn LK neurons; LHLKs) and another pair of neurons in the subesophageal ganglion (subesophageal ganglion LK neurons; SELKs) express LK in adult brain of *Drosophila*. *Lk-GAL4* also drives weak and variable expression in four pairs of neurons in the brain (approximate location of these cells is indicated by the white box; see figure S2 for an alternate preparation where these cells are weakly stained). **(B)** These four pairs of neurons do not display any LK-immunoreactivity but are positive for ITP-immunoreactivity [21] **(C)** In the ventral nerve cord (VNC), LK is expressed in eleven pairs of neurons (abdominal LK neurons; ABLKs). Seven pairs of smaller neurons in the posterior region (p) persist from the larval stages and the other four pairs (the number of pairs can vary between individuals) of larger neurons in the anterior region (a) are adult-specific [48]. **(D)** A schematic depiction of LK-expressing neurons in the brain and VNC of adult *Drosophila*. T1 – T3, thoracic neuromeres. **(E)** *Lk-GAL4* also drives ectopic expression in the salivary glands of adult *Drosophila*. **Note:** *JFRC29-10xUAS-IVS-myr::GFP-p10* was utilized in **A**, **C** and **E** whereas *UAS-mcd8-GFP* was utilized in **B**.

### DH44 is expressed in the Drosophila brain and ventral nerve cord

DH44 expression in *Drosophila* has also been examined and mapped previously [33,35]. However, different GAL4 lines result in differing expression patterns. Thus, we validated the expression of two previously generated *DH44-GAL4* lines [39,40]. One of these GAL4 lines has a minimal expression pattern and drives GFP expression in only the six MNCs in the brain (data not shown) [39]. The other *DH44-GAL4* line, which was obtained from the FlyLight collection [40], resulted in a good overlap between *DH44>GFP* and DH44-immunoreactivity. This *DH44-GAL4* drives GFP expression broadly in larval (Figure S1F, G) and adult CNS (Figure 2). In larvae, predominant expression was detected in the six MNCs in the brain (Figure S1F) and seven pairs of neurosecretory cells in the VNC (Figure S1G); however, some of these cells in the VNC were not visible in both hemiganglia. In adults, expression was again detected in the six MNCs (Figure 2A) and at least four cells dorsally (Figure 2B) and at least six cells ventrally in the posterior VNC (Figure 2C). Since there was a good overlap in GFP expression and DH44-immunoreactivity, we utilized this *DH44-GAL4* for subsequent knockdown manipulations.

**Figure 2:**
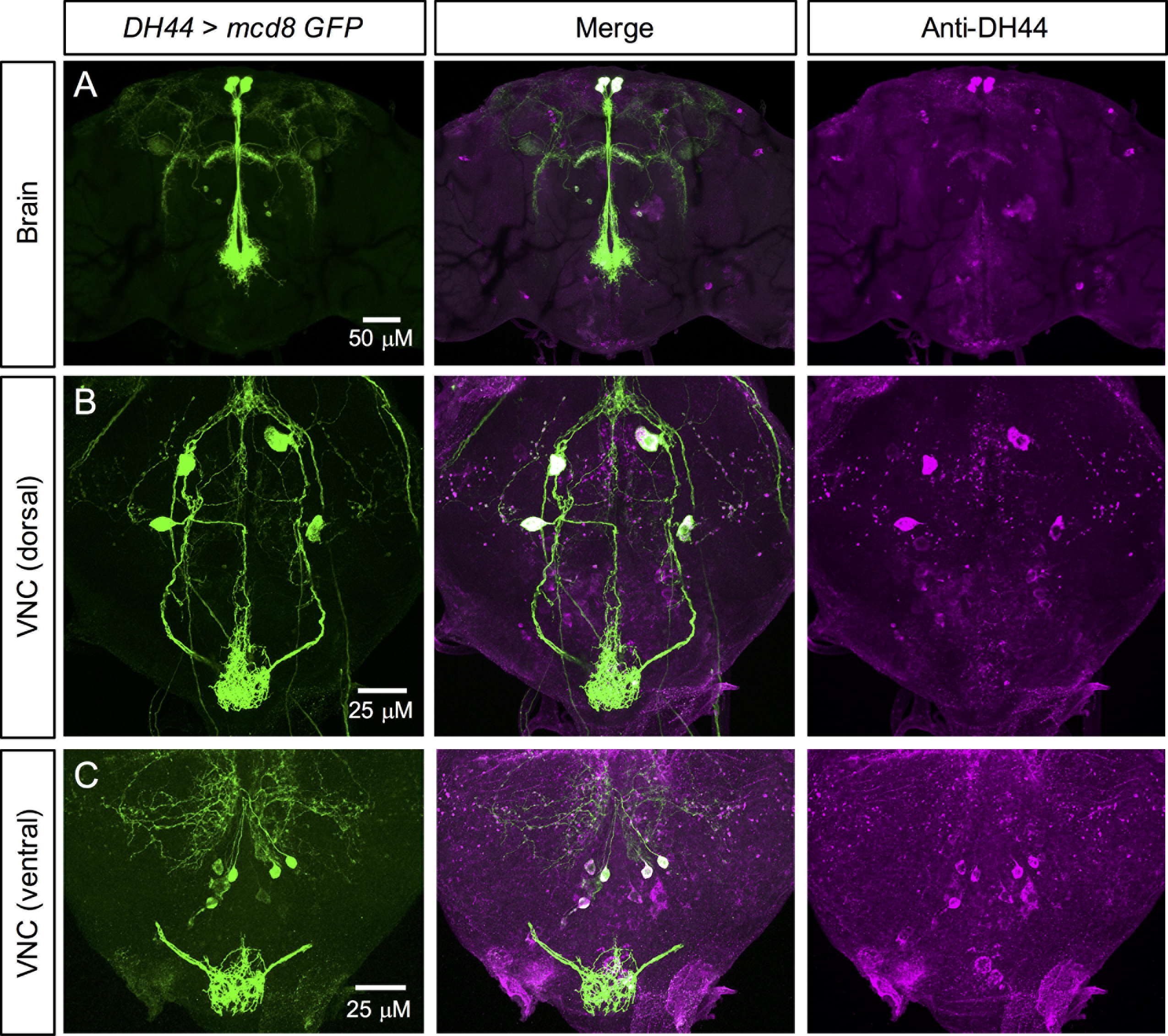
DH44 expression in the adult *Drosophila* CNS. **(A)** DH44 is expressed in three pairs of median neurosecretory cells (MNCs) in pars intercerebralis of adult *Drosophila*. Antiserum to *Drosophila* DH44 labels the same neurons identified by *DH44-GAL4*-driven GFP. **(B)** In the dorsal region of the ventral nerve cord (VNC), DH44 is expressed in two pairs of neurons. **(C)** In the ventral region of the VNC, DH44 is expressed in at least three pairs of neurons. In both B and C, there are some neurons that display DH44-immunoreactivity but do not express GFP.

### LK and DH44 are co-expressed in larval and adult ABLK neurons

The functional overlap between LK and DH44 signaling systems mentioned earlier, coupled with the expression of both LK and DH44 in the VNC neurons, prompted us to examine if these two neuropeptides are co-expressed in subsets of neurons. Our expression data show that there is no overlap in LK and DH44 expression in the brain of larvae (Figure S3A, C and Figure 4A) or adults (Figure 3A, C and Figure 4B), but these neuropeptides are co-expressed in the larval (Figure S3B, D and Figure 4A) and several adult ABLKs (Figure 3B, D and Figure 4B). Interestingly, all the larval ABLKs co-express DH44, but in adults only four to eight ABLKs express DH44 (Figure 4). Furthermore, the majority of these DH44 expressing ABLKs in the adults appear to be the ones that are generated during postembryonic neurogenesis. However, we cannot rule out the possibility that DH44 is present with a very low expression in other ABLKs.

**Figure 3:**
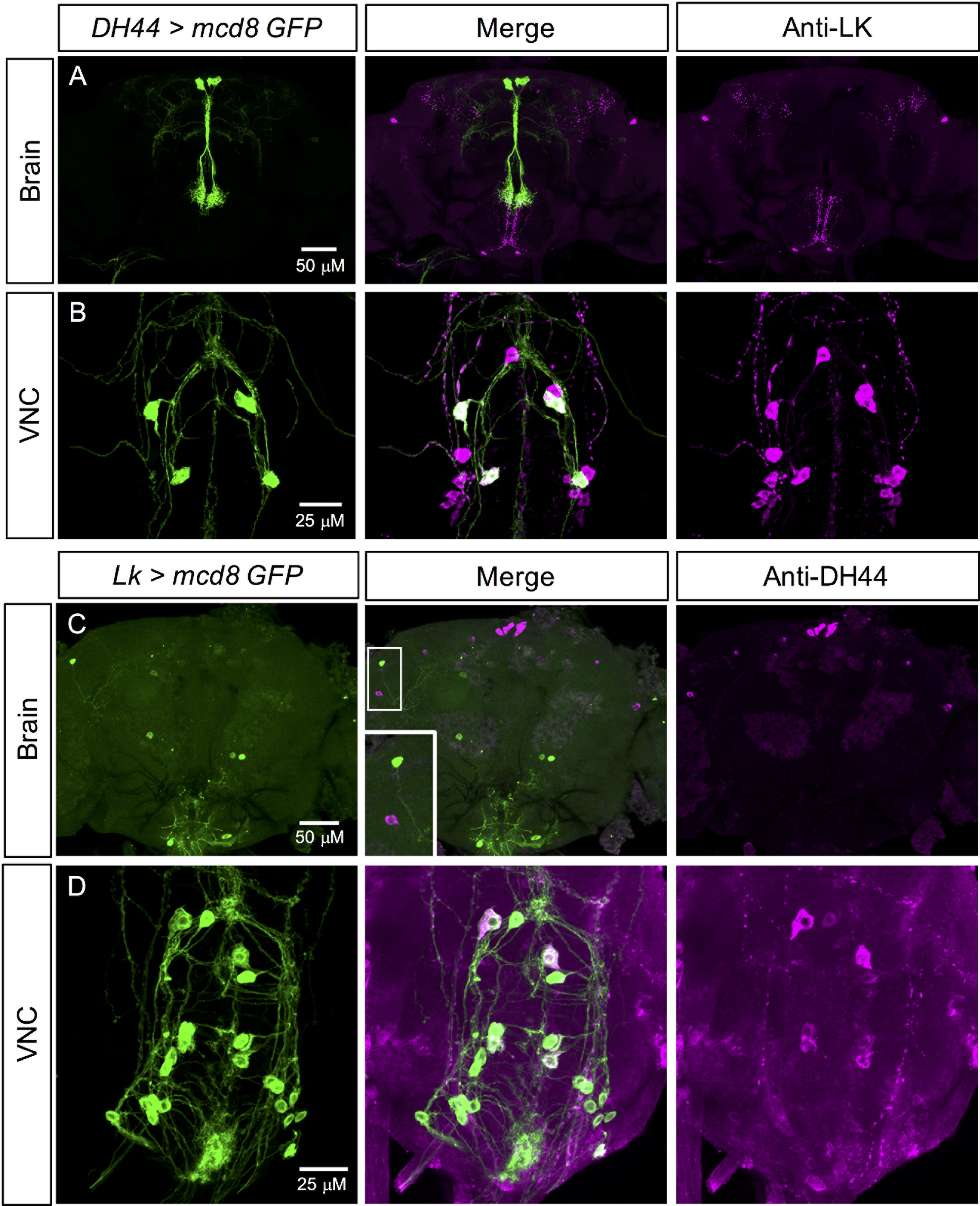
LK and DH44 are coexpressed in the ventral nerve cord, but not in the brain of adult *Drosophila*. **(A)** *DH44-GAL4* driven GFP is not colocalized with LK-immunoreactivity in the adult brain. **(B)** *DH44-GAL4* driven GFP is colocalized with LK-immunoreactivity in a subset of the abdominal LK neurons (ABLKs) in the ventral nerve cord (VNC) **(C)** *Lk-GAL4* driven GFP is not colocalized with DH44-immunoreactivity in the adult brain. **(D)** *Lk-GAL4* driven GFP is colocalized with DH44-immunoreactivity in a subset of ABLKs in the adult VNC.

**Figure 4:**
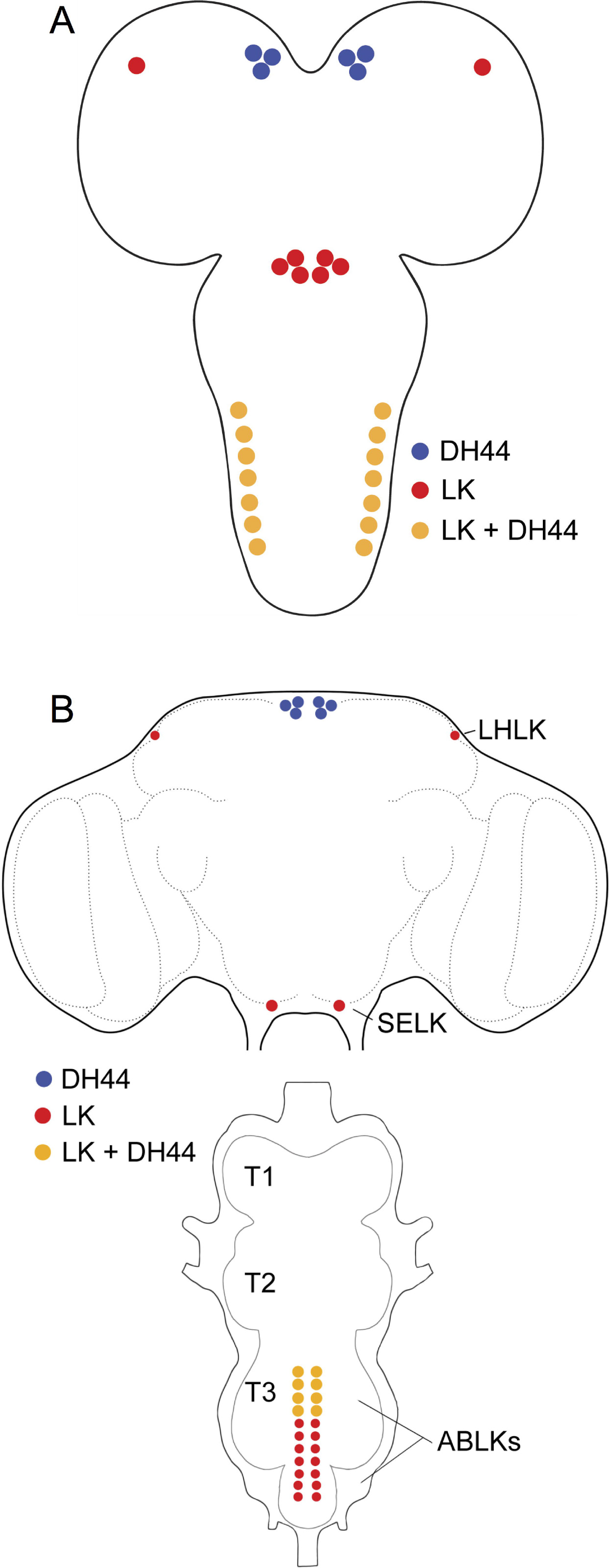
Schematics of LK- and DH44-expressing neurons in the larval and adult CNS of *Drosophila.* **(A)** A schematic of the larval CNS showing the location of neurons expressing LK, DH44 or both LK and DH44. **(B)** A schematic of the adult CNS showing the location of neurons expressing LK, DH44 or both LK and DH44. LHLK, lateral horn LK neuron; SELK, subesophageal ganglion LK neuron; ABLK, abdominal LK neuron, T1 – T3, thoracic neuromeres.

### Knockdown of Lk with Lk-GAL4 impacts stress response and water content

Having shown that LK and DH44 are co-expressed in ABLKs, it now becomes apparent that the previous studies employing genetic ablation, and activation or inactivation of LK neurons could be confounded by effects of diminishing signaling with two colocalized peptides [25,24,23]. Thus, it is not only timely to dissect the behavioral phenotypes from these previous studies using RNAi-based knockdown, but we can also study the specific functions of ABLKs using intersectional crosses. Consequently, we utilized *Lk-GAL4* and *DH44-GAL4* to knock down both *Lk* and *DH44* and assayed for effects on stress tolerance, feeding and water content. As controls in all experiments we used the parental GAL4 and UAS lines (in *w*^*1118*^ background) crossed to *w*^*1118*^ flies.

*Lk-GAL4* driven *Lk-RNAi* results in a significant decrease in LK-immunoreactivity in both the brain (Figure 5A, C) and VNC (Figure 5B). These flies with *Lk* knockdown display increased survival under desiccation (Figure 6A), starvation (Figure 6B), and ionic stress (Figure 6C), but there is no difference in chill coma recovery between experimental and control flies (Figure 6D). Moreover, food intake in CAFE assay is not affected by *Lk* knockdown (Figure 6E) (see also [22]), but these flies retain more water (Figure 6F) as demonstrated earlier [50,24]. Previous work had shown that inactivation of LK neurons resulted in increased survival under desiccation, but had no impact on starvation resistance [24]. Furthermore, in that study, both the activation and inactivation of LK neurons caused the flies to feed less in CAFE assay. This was in contrast to another study employing LK and LKR mutants, which did not find altered overall food intake, but rather an increase in meal size [22]. Hence, our data on desiccation are in agreement with previous findings and clarifies the discrepancies in previous results on food intake as possibly caused by the presence of another neuroactive compound in the LK neurons that affects feeding. Data are summarized in Table3.

**Figure 5:**
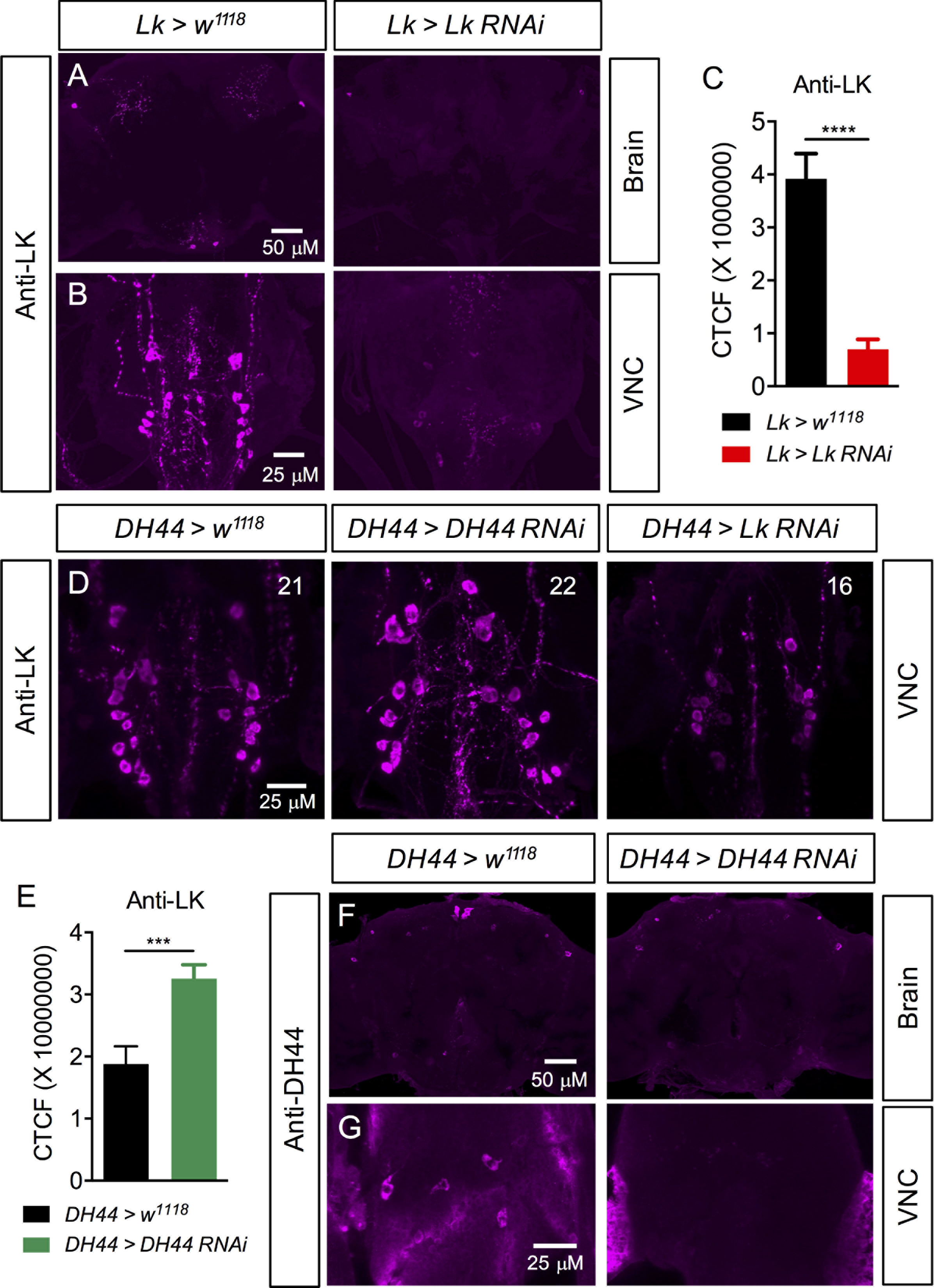
*Lk-* and *DH44-RNAi* knockdown efficiency was tested using immunolabelling. **(A, B)** Knock down of *Lk* with *Lk-GAL4* driven *Lk-RNAi* causes a significant decrease in LK-immunoreactivity in the adult brain and ventral nerve cord (VNC). (C) Fluorescence intensity measurement of lateral horn LK neurons shows a significant decrease in LK-immunoreactivity in *Lk* knock down flies compared to control flies. (*p < 0.0001, as assessed by unpaired *t* test). CTCF, corrected total cell fluorescence. **(D)** *DH44-GAL4* driven *Lk-RNAi* causes a significant decrease in LK-immunoreactivity in the adult VNC as determined by the number of immunoreactive neurons (the average number of neurons is indicated in each panel; see figure S6) that could be detected. Whereas, *DH44-GAL4* driven *DH44-RNAi* causes a significant increase in LK-immunoreactivity in adult ABLKs. (*p < 0.001, as assessed by unpaired *t* test). **(E)** and a complete abolishment of DH44-immunoreactivity in the adult brain **(F)** and VNC **(G)**.

**Figure 6:**
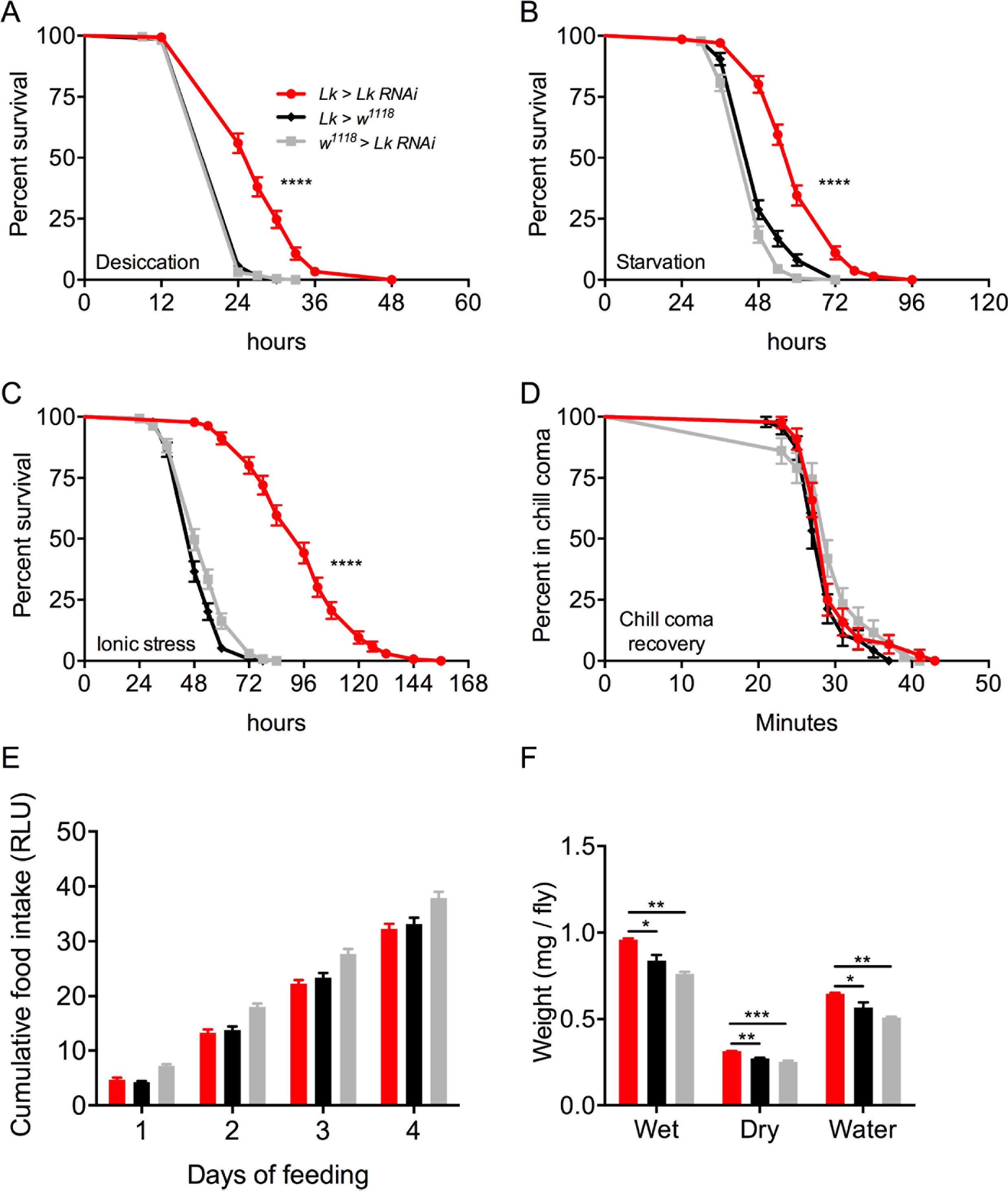
Knockdown of *Lk* using *Lk-GAL4* impacts stress resistance and water content of *Drosophila*. *Lk-GAL4* driven *Lk* knock down results in a significant increase in survival compared to control flies under **(A)** desiccation, **(B)** starvation and **(C)** ionic stress (artificial food supplemented with 4% NaCl). Data are presented in survival curves and the error bars represent standard error (**** p < 0.0001, as assessed by Log-rank (Mantel-Cox) test) **(D)** *Lk* knock down has no impact on chill coma recovery. **(E)** There is no significant difference (One-way ANOVA) in feeding as measured by capillary feeding (CAFE) assay between *Lk* knock down and control flies. Results are presented as cumulative food intake over four days. **(F)** Flies with *Lk* knock down have a higher wet weight and dry weight and retain more water (wet weight minus dry weight) compared to control flies. (* p < 0.05, ** p < 0.01, *** p < 0.001, as assessed by Oneway ANOVA). Legend for B-F is the same as the one in A.

### Knockdown of DH44 with Lk-GAL4 impacts stress response and feeding

We verified the efficiency of *DH44* knockdown using a ubiquitous driver (*Actin 5c-GAL4*) and a specific driver (*DH44-GAL4*). Knockdown of DH44 with both these drivers results in a significant decrease in DH44-immunoreactivity (Figure 5F, G and S4). We then utilized *Lk-GAL4* to knock down *DH44* specifically in ABLKs, which resulted in increased survival under desiccation (Figure 7A), starvation (Figure 7B), ionic stress (Figure 7C), and these flies display delayed recovery from chill coma compared to control flies (Figure 7D). Interestingly, *DH44* knockdown flies feed less in the CAFE assay (Figure 7E), but display no difference in water content compared to controls (Figure 7F). Hence, it appears that the effect of decreased feeding following LK neuron inactivation can be attributed to the presence of DH44 in ABLKs. Furthermore, LK but not DH44 has an effect on water content. Perhaps this could be due to the fact that LK is a more potent diuretic than DH44 in *Drosophila* and hence DH44 cannot fully compensate for the lack of LK [33,34]. Alternatively, LK could also impact water retention via actions on the hindgut [51,52]. Data are summarized in Table 3.

**Figure 7:**
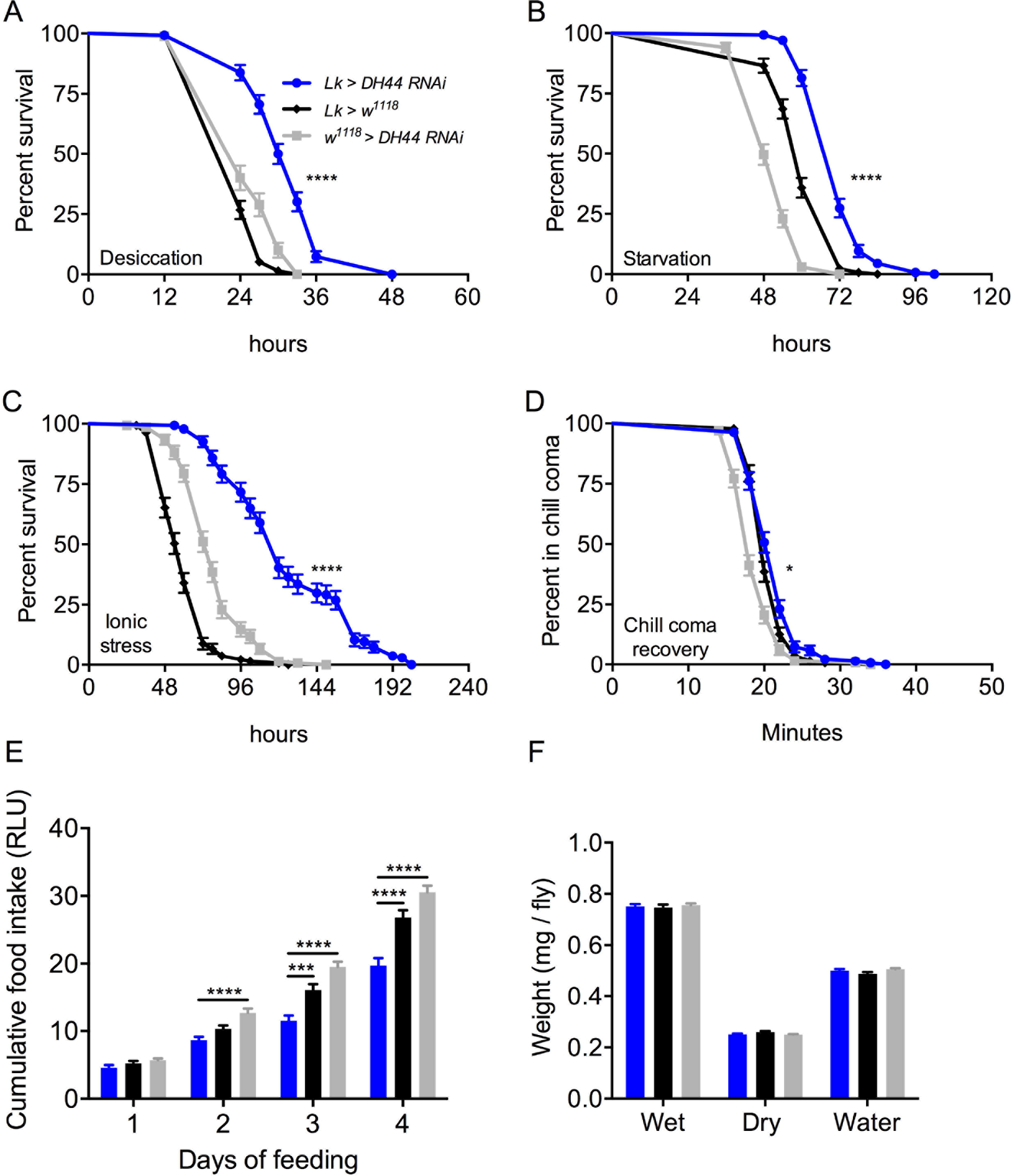
Knockdown of *DH44* using *Lk-GAL4* impacts stress resistance and feeding in *Drosophila*. *Lk-GAL4* driven *DH44* knock down results in a significant increase in survival compared to control flies under (A) desiccation, (B) starvation and (C) ionic stress (artificial food supplemented with 4% NaCl). Data are presented in survival curves and the error bars represent standard error (**** p < 0.0001, as assessed by Log-rank (Mantel-Cox) test) **(D)** *DH44* knock down causes a small delay in chill coma recovery. (* p < 0.05, as assessed by Log-rank (Mantel-Cox) test) **(E)** Flies with *DH44* knockdown feed less compared to control flies in capillary feeding (CAFE) assay. Results are presented as cumulative food intake over four days. (*** p < 0.001, **** p < 0.0001, as assessed by One-way ANOVA). **(F)** There is no significant difference in wet weight, dry weight and water content of DH44-knockdown and control flies. Legend for B-F is the same as the one in A.

### Knockdown of Lk with DH44-GAL4 impacts stress response and water content

Next, we wanted to determine the effects of knocking down *Lk* in adult-specific ABLKs. We first confirmed that *DH44-GAL4* driven *Lk-RNAi* results in an efficient knockdown in adult-specific ABLKs by counting the number of cells positive for LK-immunoreactivity (Figure 5D and S5). The average number of cells stained for LK-immunoreactivity in the control flies was 21, whereas the knockdown flies only had an average of 16 cells. Moreover, the larger adult-specific ABLKs are not labeled in the knockdown flies validating that the knockdown is efficient. Knockdown of *Lk* with *DH44-GAL4* results in increased survival during desiccation (Figure 8A), starvation (Figure 8B), ionic stress (Figure 8C), as well as a significant delay in recovery from chill coma (Figure 8D). Similar to the global *Lk* knockdown with *Lk-GAL4, Lk* knockdown in ABLKs has no effect on feeding (Figure 8E) but results in a significant increase in the water content of the flies (Figure 8F). This suggests that the effects of LK on stress response and water content could be attributed to ABLKs, and perhaps the LHLKs and SELKs of the brain play little to no part in these processes. Data are summarized in Table 3.

**Figure 8:**
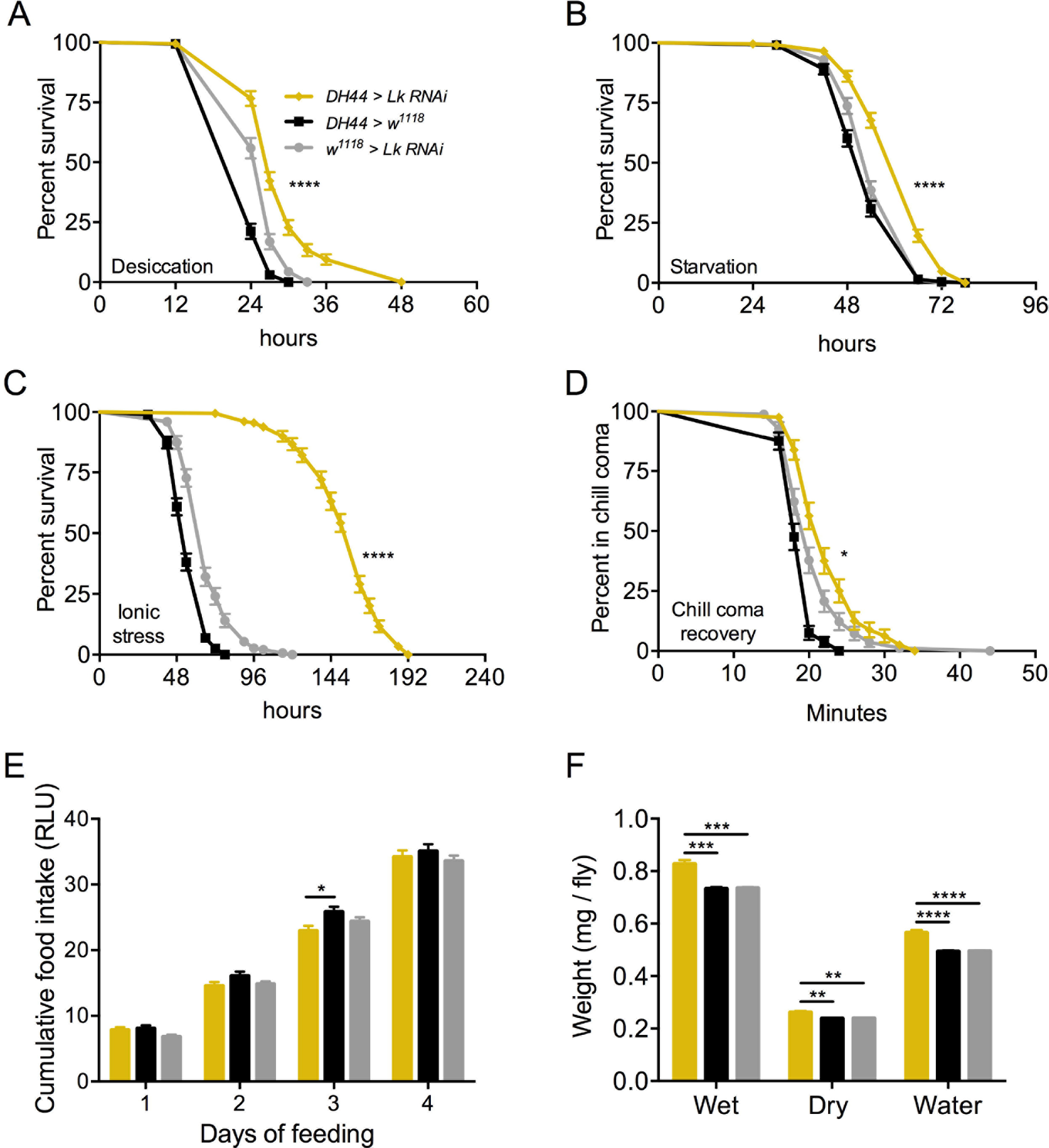
Knockdown of *Lk* using *DH44-GAL4* impacts stress resistance and water content of *Drosophila*. *DH44-GAL4* driven *Lk* knock down results in a significant increase in survival compared to control flies under **(A)** desiccation, **(B)** starvation and **(C)** ionic stress (artificial food supplemented with 4* NaCl). Data are presented in survival curves and the error bars represent standard error (**** p < 0.0001, as assessed by Log-rank (Mantel-Cox) test) **(D)** *Lk* knock down results in a delayed recovery from chill coma. (* p < 0.05, as assessed by Log-rank (Mantel-Cox) test) **(E)** There is no significant difference (One-way ANOVA) in feeding as measured by capillary feeding (CAFE) assay between *Lk* knock down and control flies. Results are presented as cumulative food intake over four days. **(F)** Flies with *Lk* knock down in ABLKs have a higher wet weight, dry weight and retain more water (wet weight minus dry weight) compared to control flies. (** p < 0.01, *** p < 0.001, **** p < 0.0001, as assessed by One-way ANOVA). Legend for B-F is the same as the one in A.

### Knockdown of DH44 with DH44-GAL4 impacts stress response and feeding

Knockdown of DH44 with DH44-GAL4 results in an efficient knockdown in the brain (Figure 5F) and VNC (Figure 5G). Staining is abolished in MNCs but not in the other cells in the brain suggesting that staining in those cells is not specific for DH44 (Figure 5F). In order to determine if there is any interaction between LK and DH44 signaling, we measured LK peptide levels in ABLKs of *DH44* knockdown flies. Interestingly, flies with *DH44* knockdown have higher LK levels suggesting that the flies may compensate for the lack of DH44 with increased LK expression (Figure 5D, E). Moreover, flies with global *DH44* knockdown display no effects on survival during desiccation (Figure S6A), but show increased resistance to starvation (Figure S6B), ionic stress (Figure S6C), and a small but significant delay in their recovery from chill coma (Figure S6D). Furthermore, flies with *DH44* knockdown display no difference in feeding (Figure S6E) and water content (Figure S6F) compared to control flies. Data are summarized in Table 3.

**Table 3.** Summary of the phenotypes obtained following different manipulations to LK and DH44 signaling. Data are compiled from Figures 6–8 and Figure S6. Notes: ↑ increase, ↓ decrease, * p < 0.05, ** p < 0.01, **** p < 0.0001.

| Assay | LK > LK Ri | DH44 > LK Ri | DH44 > DH44 Ri | LK > DH44 Ri |
| --- | --- | --- | --- | --- |
| Effect of manipulation | Global LK knockdown | LK knockdown in ABLKs | Global DH44 knockdown | DH44 knockdown in ABLKs |
| Desiccation survival | ↑**** | ↑**** | No effect | ↑**** |
| Starvation survival | ↑**** | ↑**** | ↑**** | ↑**** |
| Ionic stress survival | ↑**** | ↑**** | ↑**** | ↑**** |
| Chill coma recovery | No effect | ↑* | ↑* | ↑* |
| Feeding | No effect | No effect | No effect | ↓**** |
| Water content | ↑** | ↑**** | No effect | No effect |

### LK and DH44 act additively on Malpighian tubules to stimulate fluid secretion

Since LK and DH44 are coexpressed in ABLKs, they could potentially be coreleased into the hemolymph and result in functional interaction at the target tissue. One such site of interactions is the MTs, since both these peptides stimulate MT secretion albeit by action on different cell types and via different receptors, second messengers and ultimate targets (Cl^−^ channels for LK and V-ATPase for DH44) [33,34,53]. Hence, we were interested in examining the secretion rates by MTs and the volume of secreted fluid in the presence of either peptide alone or in the presence of both (Figure 9). Our results show that the addition of both LK and DH44 (DH44 at two concentrations) results in a secretion rate that is approximately the sum of the secretion rates obtained following the addition of each of those peptides separately (Figure 9A-D, Table 4). This additive effect is more prominent when using a higher dose of DH44 (10^−7^ M instead of 10^−9^ M) (Figure 9C, D). The amount of fluid secreted with peptide stimulation is also a reflection of these increased secretion rates (Figure 9E, F). Hence, 10^−7^ M DH44 and 10^−10^ M LK result in almost identical volumes of fluid secreted (Figure 9E), whereas a combination of both those peptides doubles the volume of fluid secreted indicating an additive response.

**Figure 9:**
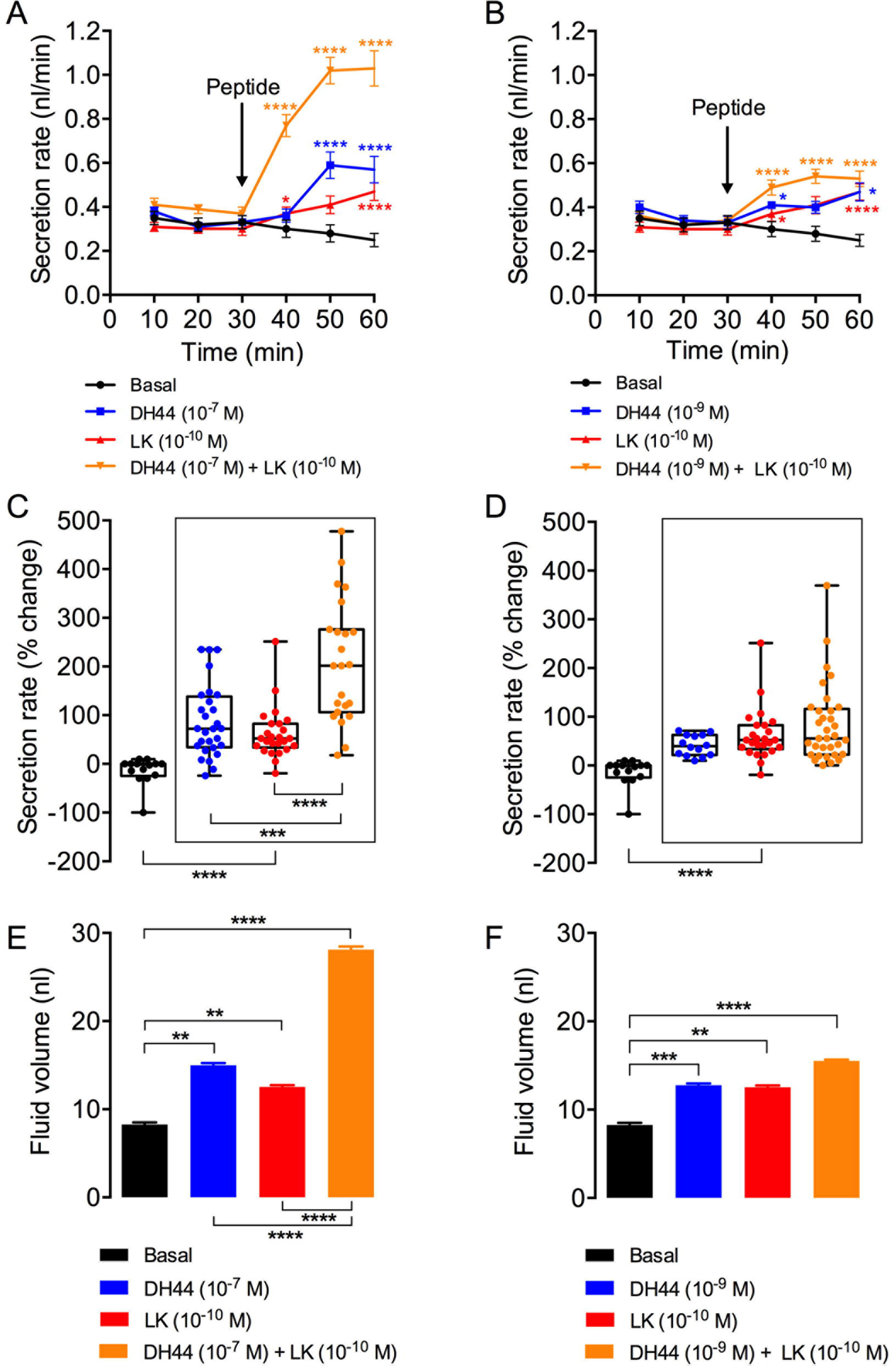
LK and DH44 peptide application results in an additive response on fluid secretion by Malpighian tubules (MTs) *ex vivo.* **(A)** Secretion rates of MTs incubated with 10^−7^ M DH44 (*n* = 28), 10^−10^ M LK (*n* = 25), a combination of both 10^−7^ M DH44 and 10^−10^ M LK (*n* = 23), or no treatment/basal (*n* = 14). **(B)** Secretion rates of MTs incubated with 10^−9^ M DH44 (*n* = 14), 10^−10^ M LK (*n* − 25), a combination of both 10^−9^ M DH44 and 10^−10^ M LK (*n* − 31), or no treatment/basal (*n* − 13). For both A and B, secretion rates were measured at 10 min intervals for 30 min before and after the addition of peptide (indicated with an arrow). Asterisk indicates significantly different secretion rate compared to basal secretion rate (secretion rate prior to the addition of peptide. For further statistics see Table 4. **(C, D)** Change (%) in secretion determined by comparing the secretion rate over the first 30 min to the maximum secretion rate following peptide application. The legend and sample size for C and D are the same as the one in A and B, respectively. **(E, F)** Total fluid secreted for 30 min following peptide application or no treatment (basal). Note that the amount of total fluid secreted following the addition of both LK (10^−10^ M) and DH44 (10^−7^ M) is a sum of the total fluid secreted following the addition of each of those peptides separately. (* p < 0.05, ** p < 0.01, *** p < 0.001, **** p < 0.0001; Mann-Whitney U test).

**Table 4:**
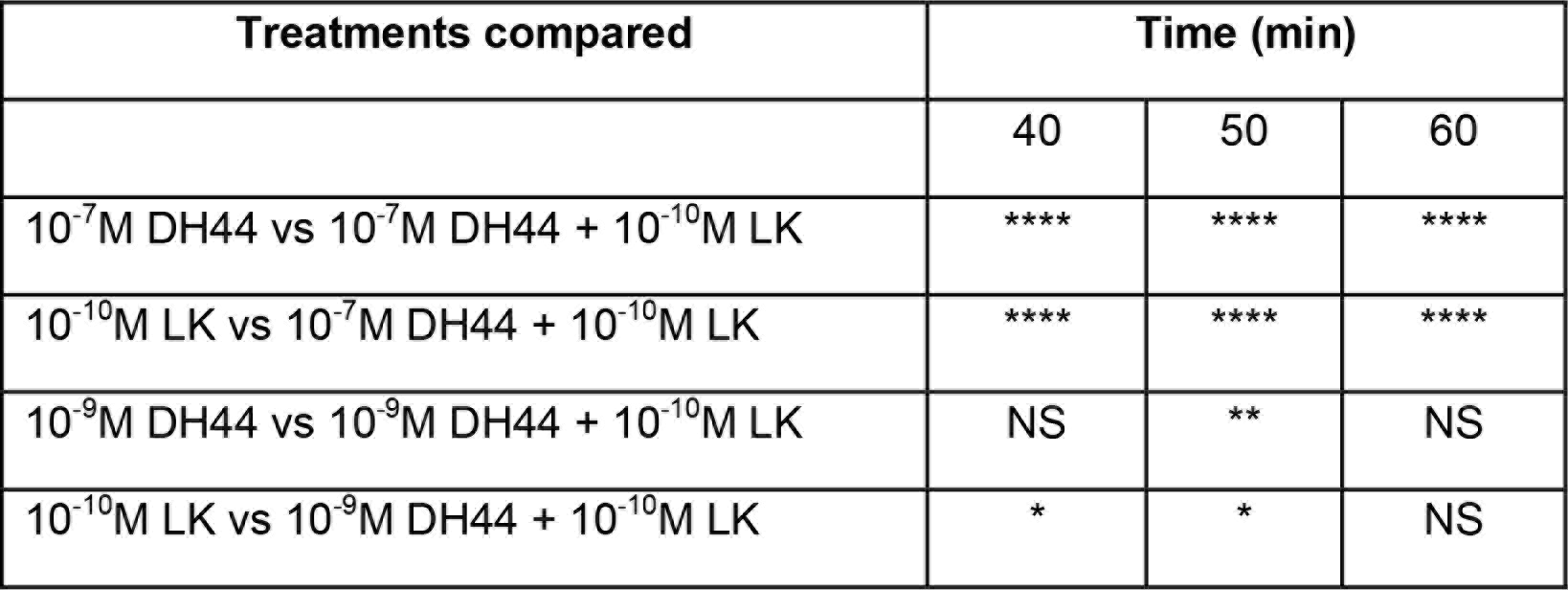
Comparison of secretion rates between various treatments and time points presented in Figure 9A and B. (NS = not significant, * p < 0.05, ** p < 0.01, **** p < 0.0001; Mann-Whitney U test).

| Treatments compared | Time (min) |  |  |
| --- | --- | --- | --- |
|  | 40 | 50 | 60 |
| $10^{-7}$M DH44 vs $10^{-7}$M DH44 + $10^{-10}$M LK | **** | **** | **** |
| $10^{-10}$M LK vs $10^{-7}$M DH44 + $10^{-10}$M LK | **** | **** | **** |
| $10^{-9}$M DH44 vs $10^{-9}$M DH44 + $10^{-10}$M LK | NS | ** | NS |
| $10^{-10}$M LK vs $10^{-9}$M DH44 + $10^{-10}$M LK | * | * | NS |

### Knockdown of Lk in ABLKs does not influence LK stimulated Malpighian tubule secretion

Previous studies have shown that knockdown of peptides could influence the expression of their receptors and vice versa (see [43]). We wanted to determine whether knockdown of *Lk* in ABLKs, the only source of hormonal LK, affects the expression of LKR in MTs, thus influencing LK-stimulated secretion by MTs. Our results indicate that there is no significant difference in LK-stimulated (10^−9^M and 10^−10^M) secretion rates of MTs isolated from *DH44* > *Lk RNAi* and control flies (Figure S7). This is similar to previous work where DH44 (10^−7^M) secretion rates were similar in tubules isolated from *DH44* > *DH44 RNAi* and control flies [23]. These results are in agreement with the *in vivo* experiments where flies with *Lk* knockdown display increased survival under desiccation.

## Discussion

Our study reveals that a portion of the LK expressing neurosecretory cells (ABLKs) in abdominal ganglia coexpress DH44, similar to earlier findings in the moth *Manduca sexta* [54], the locust *Locusta migratoria* [55] and blood sucking bug *Rhodnius prolixus* [56]. Colocalization of these peptides in multiple insect orders, including basal orders, suggests that this colocalization and the subsequent functional interaction between these signaling systems evolved early on during insect evolution. Since ABLKs are the sole neurons producing both peptides in *Drosophila* we were able to use GAL4 lines to knock down each of the two peptides in these cells only and thereby isolate the contribution of the ABLKs to physiology. This enabled us to establish that these neuroendocrine cells are sufficient for many of the functions assigned to DH44 and LK and therefore these functions are hormonally mediated. In contrast, earlier studies were based upon altering peptide levels or activity in entire populations of DH44 and LK neurons [22–26,6]. Also, we showed here that the *LK-GAL4* driver includes salivary glands and a set of ectopic brain cells (ipc-1) that do not express LK, but another peptide ITP. The ipc-1 neurons produce sNPF and tachykinin in addition to ITP and have been found to regulate stress responses [46]. This means that in earlier studies, where the *LK-GAL4* line was used to inactivate or activate neurons (see e.g. [24,25]), additional phenotypes are likely to have arisen. Using our approach, where we target only ABLK neurons, we find that both *DH44-RNAi* and *Lk-RNAi* in these cells increases resistance to desiccation, starvation and ionic stress. This suggests that diminishing the release of these two peptides from ABLKs is sufficient for this phenotype to occur. However, food intake is not affected by LK-knockdown in ABLKs, whereas DH44 knockdown diminishes feeding, and conversely knockdown of LK in ABLKs result in increased body water content, that is not seen after *DH44-RNAi*. Thus, the two colocalized peptides appear to display similar systemic actions, but differ with respect to feeding and water retention. When knocking down LK in all LK neurons we obtained a very similar set of effects as when we targeted only the ABLKs, indicating that in the assays we performed in our study, the other two sets of LK neurons (LHLK and SELK) played a minimal role.

Interestingly, knockdown of DH44 in ABLKs increases resistance to desiccation and decreases feeding, but we failed to see these effects when we diminish DH44 in all DH44 neurons. This is consistent with previous work where inactivation or activation of DH44 neurons had no effect on food intake [37]. Perhaps, the effects seen following ABLK manipulations could be compensated by action of the six DH44-expressing MNCs in the brain. Similarly, reduction of LK in ABLKs causes a slight increase in time of recovery from chill-coma, but this is not noted after global knockdown of LK. This minor difference could possibly be attributed to the strength of the two GAL4 driver lines used and, thus, the efficiency of LK knockdown in ABLKs.

We also demonstrated that DH44 and LK have additive effects on fluid secretion in MTs. It is likely that these two colocalized peptides are released together and act on the MTs where they target different cell types, receptors, signaling systems and effectors in order to regulate fluid secretion [33,34]. The action of these peptides on the MTs may also in part be responsible for the regulation of stress responses seen in our assays, as shown earlier for CAPA peptide and DH44 [57,23]. It is, however, not clear whether the altered food intake and water retention after DH44 and LK knockdown, respectively, are direct actions on target tissues or indirect effects caused by altered water and ion regulation in the fly.

Not only do the ABLKs produce two diuretic hormones, they also seem to be under tight neuronal and hormonal control. Receptors for several neurotransmitters and peptides have been identified on these cells in adults: the serotonin receptor 5-HT1B, LK receptor (LkR) and the insulin receptor, dInR [24,29]. Knockdown of the 5-HT1B receptor in ABLK neurons diminished LK expression, increased desiccation resistance, and diminished food intake, but manipulations of dInR expression in these cells generated no changes in physiology in the tests performed [24]. In larvae, all ABLKs colocalize LK and DH44, and several receptors have been detected in addition to 5-HT1B [24,30] and dInR [29], namely RYamide receptor [32], SIFamide receptor [31], and the ecdysis-triggering hormone (ETH) receptor, ETHR-A [28]. However, the expression of these receptors on adult ABLKs has so far not been investigated. Interestingly the functions of ABLKs in larvae, studied so far, seem to be primarily related to regulating muscle activity and ecdysis motor patterns. The 5-HT-1B receptor on ABLKs was shown to modulate locomotor turning behavior [30], whereas ETH mediated activation of ETHR-A on ABLKs initiates the pre-ecdysis motor activity [27,28]. In this context it is worth noting that during metamorphosis 6-8 novel ABLKs differentiate anteriorly in the abdominal ganglia [48,29], and these are the ones that display the strongest expression of DH44. In adult flies the ABLKs are neurosecretory cells with restricted arborizations in the CNS, but widespread axon terminations along the abdominal body wall and in the lateral heart nerves, whereas in larvae the same cells send axons that terminate on segmental abdominal muscles, muscle 8 [20]. It is not yet known whether larval ABLKs are involved in the regulation of diuresis and other related physiological functions *in vivo*, but certainly larval functions in locomotion and ecdysis behavior are specific to that developmental stage. Thus, it seems that there is a developmental switch of function in this set of peptidergic neuroendocrine cells.

In summary, we show that a set of abdominal neuroendocrine cells, ABLKs, coexpressing DH44 and LK are sufficient for regulation of resistance to desiccation, starvation and ionic stress, as well as modulating feeding and water content in the body. These ABLKs represent a subset of neurons that express DH44 and LK, and the functions of the remaining neurons have yet to be determined.

## Author contributions

M.Z., S.A.D. and D.R.N.: designed the research; M.Z. and R.M.: performed experiments and analyzed data; M.Z. and D.R.N.: wrote the manuscript with input from the other authors; D.R.N. and S.A.D.: obtained funding; D.R.N.: supervised the study.

## Acknowledgements

The authors would like to acknowledge Dr. Yiting Liu and Dr. Olga Kubrak for very helpful advice and valuable discussions. We are grateful to the Bloomington *Drosophila* Stock Center, the Vienna *Drosophila* Resource Center and Drs. Jan A. Veenstra, Pilar Herrero, Young Joon Kim and Michael Texada for providing flies and reagents. Stina Hoglund and the Imaging Facility at Stockholm University (IFSU) are acknowledged for maintenance of the confocal microscopes. This work was supported by a European Commission Horizon 2020, Research and Innovation Grant 634361.

## Supplementary files

**Figure S1: LK and DH44 expression in the larval *Drosophila* CNS. (A)** *Lk-GAL4* drives expression in five pairs of neurons in the brain; however, four of these pairs do not display any LK-immunoreactivity [21]. **(B)** These four pairs of neurons display ITP-immunoreactivity. **(C)** Three pairs of neurons in the subesophageal ganglion express *Lk* in larval *Drosophila*. **(D)** Seven pairs of neurons in the larval ventral nerve cord (VNC) express *Lk*. **(E)** A schematic of LK-expressing neurons in the larval brain and VNC of *Drosophila*. Neurons displaying LK-immunoreactivity are labeled in red and neurons displaying ITP-immunoreactivity are labeled in black. **(F)** *DH44-GAL4* driven GFP and DH44-immunoreactivity is present in three pairs of median neurosecretory cells in the larval brain. **(G)** DH44 is expressed in several neurons, with strong expression seen in seven pairs of neurosecretory cells in the larval VNC. In both F and G, there are some neurons that contain GFP but do not contain DH44-immunoreactivity.

**Figure S2: LK expression in adult *Drosophila* brain**. *Lk-GAL4* drives weak GFP expression in four pairs on neurons in the adult brain. The location of these cells is indicated by white boxes.

**Figure S3: LK and DH44 are coexpressed in neurons of the ventral nerve cord but not in the brain of larval *Drosophila*. (A)** *DH44-GAL4* driven GFP is not colocalized with LK-immunoreactivity in the larval brain. **(B)** *DH44-GAL4* driven GFP is colocalized with LK-immunoreactivity in all seven pairs of abdominal LK neurons (ABLKs) in the ventral nerve cord (VNC) **(C)** *Lk-GAL4* driven GFP is not colocalized with DH44-immunoreactivity in the larval brain. **(D)** *Lk-GAL4* driven GFP is colocalized with DH44-immunoreactivity in ABLKs in the larval VNC.

**Figure S4: Knockdown of *DH44* using *Actin5c-GAL4***. *Actin5c-GAL4* driven *DH44* knockdown results in a complete abolishment of DH44-immunoreactivity in the six neurons in pars intercerebralis of adult *Drosophila*.

**Figure S5: Number of LK-immunoreactive neurons in the VNC following knockdown of *DH44* and *Lk* using *DH44-GAL4***. *Lk* knockdown but not *DH44* knockdown causes a significant decrease in the number of LK-immunoreactive neurons that could be detected in the adult VNC. (**** p < 0.0001, as assessed by One-way ANOVA).

**Figure S6: Knockdown of *DH44* using *DH44-GAL4* impacts stress resistance and feeding in *Drosophila***. *DH44-GAL4* driven *DH44* knock down results in a significant increase in survival compared to control flies under **(A)** desiccation (compared to the GAL4 control), **(B)** starvation and **(C)** ionic stress (artificial food supplemented with 4% NaCl). Data are presented in survival curves and the error bars represent standard error (**** p < 0.0001, as assessed by Log-rank (Mantel-Cox) test) **(D)** *Lk* knock down results in a delayed recovery from chill coma. (* p < 0.05, as assessed by Log-rank (Mantel-Cox) test) **(E)** There is a significant decrease (One-way ANOVA) in feeding as measured by capillary feeding (CAFE) assay in *DH44* knock down flies (compared to the GAL4 control). Results are presented as cumulative food intake over four days. **(F)** There is no significant difference in wet weight, dry weight and water content of *DH44*-knockdown and control flies. Legend for B-F is the same as the one in A.

**Figure S7: Knockdown of *Lk* in ABLKs with *DH44-GAL4* does not influence LK-stimulated Malpighian tubule secretion *ex vivo*. (A)** Secretion rates of 10^−9^ M LK stimulated MTs isolated from *DH44* > *w*^1118^ (*n* = 14) or *DH44* > *Lk RNAi* flies (*n* = 25). **(B)** Secretion rates of 10^−10^ M LK stimulated MTs isolated from *DH44* > *w*^1118^ (*n* = 10) or *DH44* > *Lk RNAi* flies (*n* = 12). For both A and B, secretion rates were measured at 10 min intervals for 30 min before and after the addition of peptide (indicated with an arrow). **(C, D)** Change (%) in secretion determined by comparing the secretion rate over the first 30 min to the maximum secretion rate following peptide application. The legend and sample size for C and D are the same as the one in A and B, respectively. Asterisk indicates significantly different secretion rate compared to basal secretion rate (secretion rate prior to the addition of peptide). (NS = not significant, * p < 0.05, ** p < 0.01, *** p < 0.001; Mann-Whitney U test)

